# Highly Sensitive Detection of *Campylobacter spp.* in Chicken Meat using a Silica Nanoparticle Enhanced Dot Blot DNA Biosensor

**DOI:** 10.1101/2020.07.03.185827

**Authors:** Priya Vizzini, Marisa Manzano, Carole Farre, Thierry Meylheuc, Carole Chaix, Nalini Ramarao, Jasmina Vidic

**Affiliations:** Université Paris-Saclay, INRAE, AgroParisTech, Micalis Institute, 78350, Jouy-en-Josas, France; Dipartimento di Scienze AgroAlimentari, Ambientali e Animali, Università di Udine, Italy; Institut des Sciences Analytiques, UMR 5280, CNRS-UCBL, Université de Lyon, Lyon, France

**Keywords:** *Campylobacter*, DNA dot blot, Si-nanoparticles, Food safety, Multiplex bacterial detection

## Abstract

Paper-based DNA biosensors are powerful tools in point-of-care diagnostics since they are affordable, portable, user-friendly, rapid and robust. However, their sensitivity is not always as high as required to enable DNA quantification. To improve the response of standard dot blots, we have applied a new enhancement strategy that increases the sensitivity of assays based on the use of biotinylated silica-nanoparticles (biotin-Si-NPs). After immobilization of a genomic *Campylobacter* DNA onto a paper membrane, and addition of a biotinylated-DNA detection probe, hybridization was evidenced using streptavidin-conjugated to horseradish peroxidase (HRP) in the presence of luminol and H_2_O_2_. Replacement of the single biotin by the biotin-Si-NPs boosted on average a 30 fold chemiluminescent read-out of the biosensor. Characterization of biotin-Si-NPs onto a paper with immobilized DNA was done using a scanning electron microscope. A limit of detection of 3 pg/μL of DNA, similar to the available qPCR kits, is achieved, but it is cheaper, easier and avoids inhibition of DNA polymerase by molecules from the food matrices. We demonstrated that the new dot blot coupled to biotin-Si-NPs successfully detected *Campylobacter* from naturally contaminated chicken meat, without needing a PCR step. Hence, such an enhanced dot blot paves the path to the development of a portable and multiplex paper based platform for point-of-care screening of chicken carcasses for *Campylobacter*.

**Graphical abstract:** 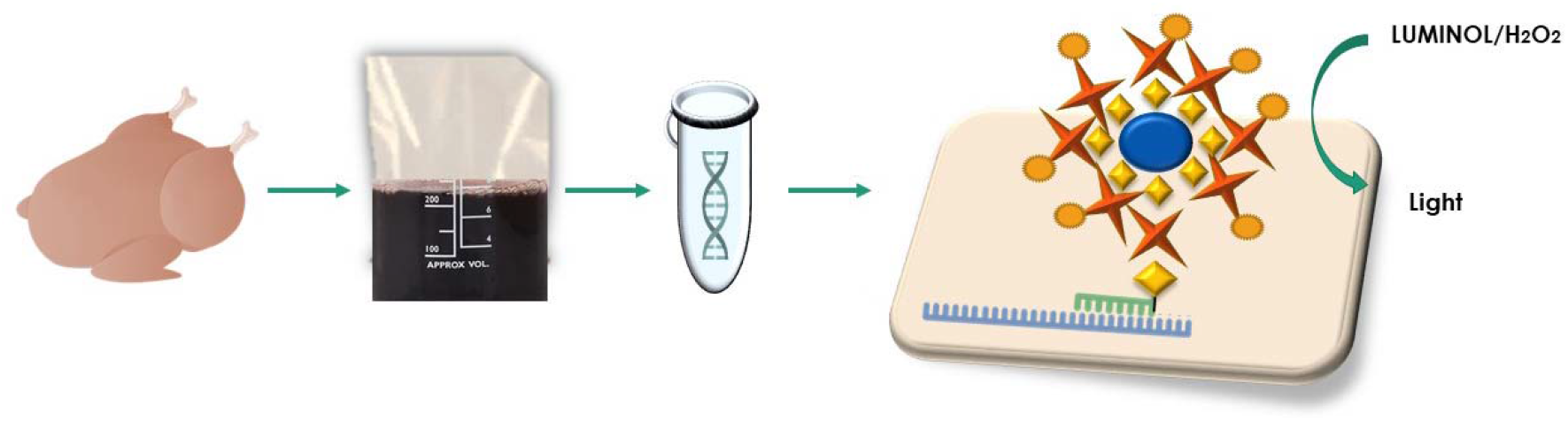

## 1. Introduction

*Campylobacter* is considered the most common bacterial cause of human gastroenteritis in the world. The World Health Organization (WHO) estimates that 550 million people fall ill every year from gastroenteritis, of which 220 million are children (WHO 2020). This zoonosis is transmitted mostly through exposure to under-cooked poultry products (50-80% cases), and in a minor way, to contaminated milk, vegetables and water (ECDC 2018; Hermans et al. 2012; Humphrey et al. 2007). Campylobacteriosis in humans is usually a self-limiting condition involving bloody diarrhea, abdominal cramping, nausea and fever, which can last up to two weeks. In 1 % of cases, campylobacteriosis evolves to the Guillain-Barré syndrome, a severe autoimmune disease that leads to death in 2–12 % of patients, depending on their age (Scallan Walter et al. 2020). The overall economic burden of campylobacteriosis was estimated to about EUR 3 billion/year in EU (ECDC 2018), and between US$ 1.2 and 4 billion/year for the US (Batz et al. 2014). To prevent the entry of *Campylobacter* contaminated broilers into the market, the European Commission adopted a process hygiene criterion (EU 2017/1495) with the critical limit of 1000 cfu/g of broiler meat or skin and obligatory systematic screening of broiler carcasses for *Campylobacter* spp. (EU No 2017/625).

Identification and quantification of *Campylobacter* spp. rely on official, culture-based methods, and bacterial biochemical/phenotypical characterization (Vizzini et al. 2019). *Campylobacter* is highly infectious, with reported infective doses as low as 500 cells (Black et al. 1988). To ensure the detection of one cell of *Campylobacter* in 25 g of food, an enrichment step in Bolton broth for 24–48 h is needed, before isolation of the colonies by culturing (it takes about 48–72 h) onto selective agar plates incubated in chambers for microaerophilic conditions. *Campylobacter* is usually present in low numbers in food samples when compared to bacteria like coliforms and Enterobacteriaceae making its isolation on common agar media difficult. The official ISO 10272-1:2006 method for *Campylobacter* detection may provide false negative results because of the possibility *Campylobacter* death during handling. Furthermore, *Campylobacter* can enter in a viable but not cultivable (VBNC) status in food matrices, making its detection based on culturing impossible (Vidic et al. 2017; Vizzini et al. 2019). Identification of *Campylobacter* by the optical microscope is not easy as bacteria can change their spiral distinctive shape into spherical or coccoid. Moreover, the current problem of official methods is the use of a not enough selective medium, and emergence of bacterial resistance against the antibiotic added to the enrichment broth.

Culture-independent molecular methods, such as PCR, and real-time PCR are used as alternatives to colony growth (De Boer et al. 2015; Fontanot et al. 2014; Gosselin-Théberge et al. 2016; Liu et al. 2017; Ricke et al. 2019). However, PCR-based methods may also provide false-negative responses because of the sensitivity of DNA polymerase to inhibitors present in food matrices and enrichment broths (Schrader et al. 2012; Vidic et al. 2019). Next generation sequencing (NGS) which enables sequencing of the entire bacterial genome in a relatively short time has started to be used as a tool for the identification of infectious bacteria. However, NGS can hardly be routinely used at farms and slaughterhouse because it requires special equipment and highly trained personnel for data interpretation (Gosiewski et al. 2017).

Biosensors for detection of *Campylobacter* show marked advantages over traditional methods in terms of rapidity, facility utilization and cost-effectiveness (Manzano et al. 2015; Masdor et al. 2016; Morant-Miñana and Elizalde 2015; Vidic et al. 2017; Vidic et al. 2019; Vizzini et al. 2019; Yang et al. 2013). However, today no commercial biosensor is available to detect *Campylobacter* in food matrices, mainly due to the difficulty of attaining a high sensitivity.

Due to significant improvement in biosensor technology over the last two decades, applications of paper-based sensors in pathogen detection are increasing. Such devices seem to meet ASSURED criteria (Affordable, Sensitive, Specific, User□friendly, Rapid and robust, Equipment free and Deliverable) recommended by the WHO for point-of-care diagnostics. Paper as a substrate has the advantage of being inexpensive, lightweight, and easily enables multiplex analysis (Dincer et al. 2019). Various paper-based methods to detect foodborne pathogens have been reported including enzymatic-, immuno- and DNA/RNA- tests (Jokerst et al. 2012; Morales-Narváez et al. 2015; Russell et al. 2017). Tests that target nucleic acids are among the most accurate and specific although they are usually associated with a PCR step, including isothermal amplification, to allow for detection of pathogens in low titers (Trinh et al. 2020; Vidic et al. 2019).

Here, we coupled functionalized silica nanoparticles (Si-NPs) to a paper based DNA dot blot test to enable sensitive *Campylobacter* nucleic acid detection without a pre-amplification step. A highly specific DNA probe that recognizes the 16S rRNA gene of the most prevalent *Campylobacter* spp. causing infections (*C. jejuni*, *C. coli*, *C. lari*, and *C. upsaliensis*) was used in the test. Si-NPs decorated with biotin (biotin-Si-NPs) were employed to enhance the strepatavidin–HRP chemiluminescent signal read-out. Our study illustrates the proof-of-concept that the collective effect of biotin-Si-NPs could become an effective means for increasing the sensitivity of the cost-effective, specific and easy-to-perform dot blot assay.

## 2. Material and Methods

### 2.1 Materials and reagents

Streptavine-HRP, Proclin, acetonitrile, 1,8-Diazabicyclo[5.4.0]undéc-7-ène (DBU), Controlled pore glass (120-200 Mesh, CPG-3000 Å) were purchased from Sigma-Aldrich (Saint Quentin Fallavier, France). Fluorescent rhodamine B silica NPs (Si-NPs, diameter~50 nm) were supplied by Nano-H (Saint Quentin Fallavier, France). DNA phosphoramidite synthons and all DNA-synthesis reagents were purchased from Glen Research (Sterling, Virginia, USA). Phosphate buffered saline (PBS) was purchased from Dominique Dutscher (Brumath, France). Amersham Hybond-N+ nylon and Amercham Hybon-XL nitrocellulose membaranes were purchased from ThermoFisher (Illkirch, France).

All bacterial media and supplements used were from Oxoid (Milan, Italy), except for the Violet red Bile glucose (VRBG) agar and Coli ID medium that were purchased from Biomerieux (Bagno a Ripoli, Italy). Triton, SDS, NaCl, trizma-base, phenol, chloroform, bacteriological peptone and isoamyl alcohol were purchased from Sigma (Milan, Italy). A 36- mer oligonucleotide related to the 16S gene encoding for ribosomal *Campylobacter* RNA of *C. jejuni*, *C. coli*, *C. lari*, and *C. upsaliensis* (base location: 72-108) was used as a recognition element (probe named CampyP3). It is interesting to note that the probe matches in three points for each *Campylobacter* genome at 100%. CampyP3 was labeled at 5’ with biotin for chemiluminescent dot blot assay development. The probe was tested *in silico* by the OligoAnalyzer3.1 (https://eu.idtdna.com/calc/analyzer), the Amplifix software, Fast PCR 6.1 and Blast (https://blast.ncbi.nlm.nih.gov/Blast.cgi). An ssDNA sequence complementary to the CampyP3 probe, of the same length as the probe was named CP3 and used as positive control. Two ssDNA sequences of the same length as the CP3, but not complementary to CampyP3, were named PR and PE, and used as negative controls to study selectivity of the sensor through their hybridization with the probe. PR was designed by mismatching positions of nucleic acids of the CP3 sequence, while PE corresponded to the sequence of *E. coli (*accession number 527445.1) (base location: 338-376 for *E. coli),* which shows some similarities to *Campylobacter.* These sequences, reported in Table 1S, were provided by Sigma-Aldrich (Saint Quentin Fallavier, France) as lyophilized powers. All solutions were prepared using Milli-Q water.

### 2.2 Bacterial strains

Bacteria used in this study are listed in Table 2S. *Campylobacter* strains were grown under microaerophilic conditions (5% O_2_, 10 % CO_2_ and 80% N_2,_ generated with a Sachet Oxoid™ CampyGen™ 2.5 L (Oxoid, Italy) in anaerobic gas jars at 37°C for 48h, on Columbia blood agar plates. *Campylobacter* isolates, both reference strains and strain isolates from chicken samples, were subjected to Gram straining and optical microscope observations for cell morphology and motility (Brucella broth, Thermofisher, Milan, Italy) after oxidase and catalase tests.

All negative control strains were grown on their specific culture media at optimal conditions and subjected to the same tests carried out for *Campylobacter* before DNA extractions.

### 2.3 Sample collection, plate count enumeration and selective isolation

Five chickens were purchased from local supermarkets in Italy. 10 g of chicken skin was transferred into a filter-sterile stomacher bag with 40 mL saline-peptone water (8 g/L NaCl, 1 g/L bacteriological peptone) and homogenized in a Stomacher (PBI, Milan, Italy) for 90 s. Aliquots of 0.1 mL were spread for the mesophilic aerobic count on Triptone Soya Agar, and incubated at 30°C for 48 h, while 0.1 mL were spread on Agar Malt tetracycline at 30°C for 48 h to count yeast/molds. Aliquots of 1 mL were used with the pour plate method for enumeration of Enterobacteriaceae in the VRBG agar (37°C for 24 h), and coliforms and *E. coli* in Coli ID medium (37°C for 24 h). *Campylobacter* detection was performed according to the conventional method ISO 10272-1:2006. For selective bacterial isolation, 10 μL of the Bolton broth were streaked onto Skirrow agar plates, and mCCDA (modified charcoal-cefoperazone-deoxycholade) plates and incubated at 41.5 °C for 48h, under microaerophilic conditions. One colony was selected from mCCDA and streaked on two plates of Columbia blood agar. One plate was incubated at 41.5 °C for 48 h in aerobic condition and one plate at 25 °C for 48 h in microphilic condition. Growth on Skirrow served as a confirmation to proceed with the identification steps.

### 2.4 DNA extraction from pure cultures and enrichment broths

DNA was extracted from reference strains and isolates from chicken samples according to (Manzano et al. 2015). 2 mL of Bolton enrichment broth was centrifuged at 13,000 g at 4 °C for 10 min, and bacterial pellets were washed three times with PBS 1X, and subjected to extraction (Manzano et al. 2015). Extracted DNAs were rehydrated using 50 μL of sterile distilled water and treated with RNase enzyme at 37°C for 1 h. Finally, DNA was quantified using a Nanodrop™ 2000C (ThermoFisher Scientific, Milan Italy) spectrophotometer. The concentration of DNA was adjusted to 100 ng/μL using sterile distilled water.

### 2.5 Si-NP synthesis, labeling and quantification

Si-NPs was functionalized by an innovative solid-phase synthesis technology which enables functionalization of nanosized particles with DNA fragments, as reported previously (Bonnet et al. 2018; De Crozals et al. 2012). Briefly, nanoparticle immobilization onto controlled pore glass (CPG) allows a very high functionalization with a modified oligonucleotidic based linker. The linker (sequence: dT10-PEG2-dT10) was synthetized using an applied Biosystems 394 RNA/DNA synthesizer (Applied Biosystems). After grafting nanoparticles on a CPG support, they were functionalized by automated synthesis using the phosphoramidite chemistry. First, the linker, that allows a better accessibility of the functions of interest, was synthesized. Second, the biotin group was incorporated. We controlled the biotin incorporation to reach a 10% coupling yield. To do it, diluted solutions of biotin phosphoramidite (10 mM) and tetrazole in acetonitrile (45 mM) were used and the coupling time was reduced to 10 s. The biotin incorporation was monitored at 498 nm using dimethoxytrityl quantification by an UV-visible spectrophotometer. Third, biotin-Si-NPs were released from CPG by incubating the support in 1 mL of 0.1% (m/v) DBU in water-acetonitrile 1/1 (v/v). The DBU solution was stirred in a thermomixer at 22°C during 1 h before recovering. A fresh DBU solution was added to the CPG suspension every hour. The release kinetics was followed by quantifying the NP concentration in each DBU solution with an UV-visible measurement at 560 nm. Released nanoparticles were washed with milli-Q water (1x 4 mL then 3 × 2 mL) and concentrated on 30 K Amicon Ultra filter (5000 g, 10 min). Finally, the amount of dT10-PEG2-dT10-10% Biotin strands per NP was estimated using a Varian Cary 100 Bio UV-visible spectrophotometer (Agilent Technologies) and a quartz cuvette of 1 cm path length.

The amount of strands per NP was estimated using a Varian Cary 100 Bio UV-visible spectrophotometer (Agilent Technologies) and a quartz cuvette of 1 cm path length. The amount of linker grafted to nanoparticles was quantified by measuring absorbance in water (200 μL) at 260 nm and 560 nm with a microplate reader (Perkin Elmer). The nanoparticle concentration was estimated as described previously (Bonnet et al. 2018). To fit with this estimation, we approximated a molar extinction coefficient to 163200 M^−1^ cm^−1^ that considers the epsilons of the different parts of the sequence corrected by their corresponding coupling yield.

Functionalized NPs were observed under a transmission electron microscopy (TEM) using a Philips CM120 instrument operating at an accelerate voltage of 120 kV (Centre Technologique des microstructures, Lyon). Si-NPs were observed after deposition of 5 μL of diluted solution on a formvar-carbon coated copper grid followed by evaporation until dry.

### 2.6 Scanning electron microscopy

Samples were mounted on aluminum stubs (50 mm diameter) with carbon adhesive discs (Agar Scientific, Oxford Instruments SAS, Gomez-la-Ville, France) and visualized by field emission gun scanning electron microscopy (SEM FEG) as secondary and backscattered electrons images (5 kV) under high vacuum conditions with a Hitachi SU5000 instrument (Milexia, Verrières-le-Buisson, France). Sample preparation and scanning Electron Microscopy analyses were performed at the Microscopy and Imaging Platform MIMA2 (INRAE, Jouy-en-Josas, France).

### 2.7 Sample immobilization and detection procedure

Prior to immobilization onto the nylon membrane, extracted DNAs were denatured at 95 °C for 10 min, put immediately on ice and 1 μL spotted on the positively charged nylon membrane (Amersham Hybon™-XL, GE Healthcare, France), which was exposed to UV at 254 nm for 10 min to fix DNA. The membrane was then soaked in a pre-wormed hybridization buffer (0.5 M Na_2_HPO_4_, 0.5 M NaH_2_PO_4_, 10 mM EDTA, 1 % SDS, pH 7.5) at 65°C for 30 min under gentle shaking. 4 ng/μL of the denatured biotin labeled CampyP3 probe (100 ng/μL) was added to the hybridization buffer and left overnight at 65°C under gentile shaking to allow for hybridization.

Subsequently, the membrane was washed twice with 0.1 SDS, 300 mM SSC (Saline sodium citrate, Meraudex, France) for 5 min and with 75 mM SSC for 15 min. The membrane was transferred in the blocking buffer (0.1 % Tween, 0.1 % BSA, 0.03% Proclin, PBS, pH 7.4) at room temperature for 15 min to saturate the surface. The membrane was finally incubated with the blocking solution containing 0.7 μM streptavidin-HRP for 15 min at room temperature. After washing with 0.1 % SDS, 150 mM SSC at room temperature for 5 min, the membrane was incubated with 10^6^ biotin-Si-NPs/mL, PBS, pH 7.4 under gentile shaking at room temperature, for 30 min. A 0.7 μM streptavidin-HRP in blocking solution was added after washing and incubated for 30 min with shaking. The signal was then revealed using the enhanced chemiluminescent substrate for detection HPR (Thermo Scientific, France). The membrane was removed from the solution and observed under a ChemiDoc MP imaging system (Biorad, France). Detection signals were quantified using Image Lab™ software (Biorad). The normalized value of a spot intensity was calculated by (PI_0_ –PI_n_)/PI_0_, where PI_0_ and PI_n_ represent the pixel intensity obtained for detection of 0.1 ng/μL CP3 probe (standard) and the experimental sample, respectively.

## Results and discussion

### 3.1. Preparation and characterization of biotin-Si-NPs

Biotin-Si-NPs were prepared by an innovative solid-phase synthesis technology that allowed for a very high functionalization. A hydroxyl group was attached to the Si-NP surface by silanization to support the oligonucleotide linker synthesis on the NP surfaces, as previously reported (Bonnet et al. 2018; Farre et al. 2010). In the final step, biotin groups were grafted to the linker. TEM observations showed that the NP morphology was stable during synthesis (Fig. 1A). Functionalized biotin-Si-NPs had an average size of 50 ± 3 nm, as estimated from TEM images (Fig. 1B). The absorbance measurements suggested that about 500 molecules of the linker were attached to each NP, while dimethoxytrityl quantification indicated the biotin-coupling yield of 10 %, which quantified biotin to about 50 per NP (Fig. 1C).

**Figure 1.**
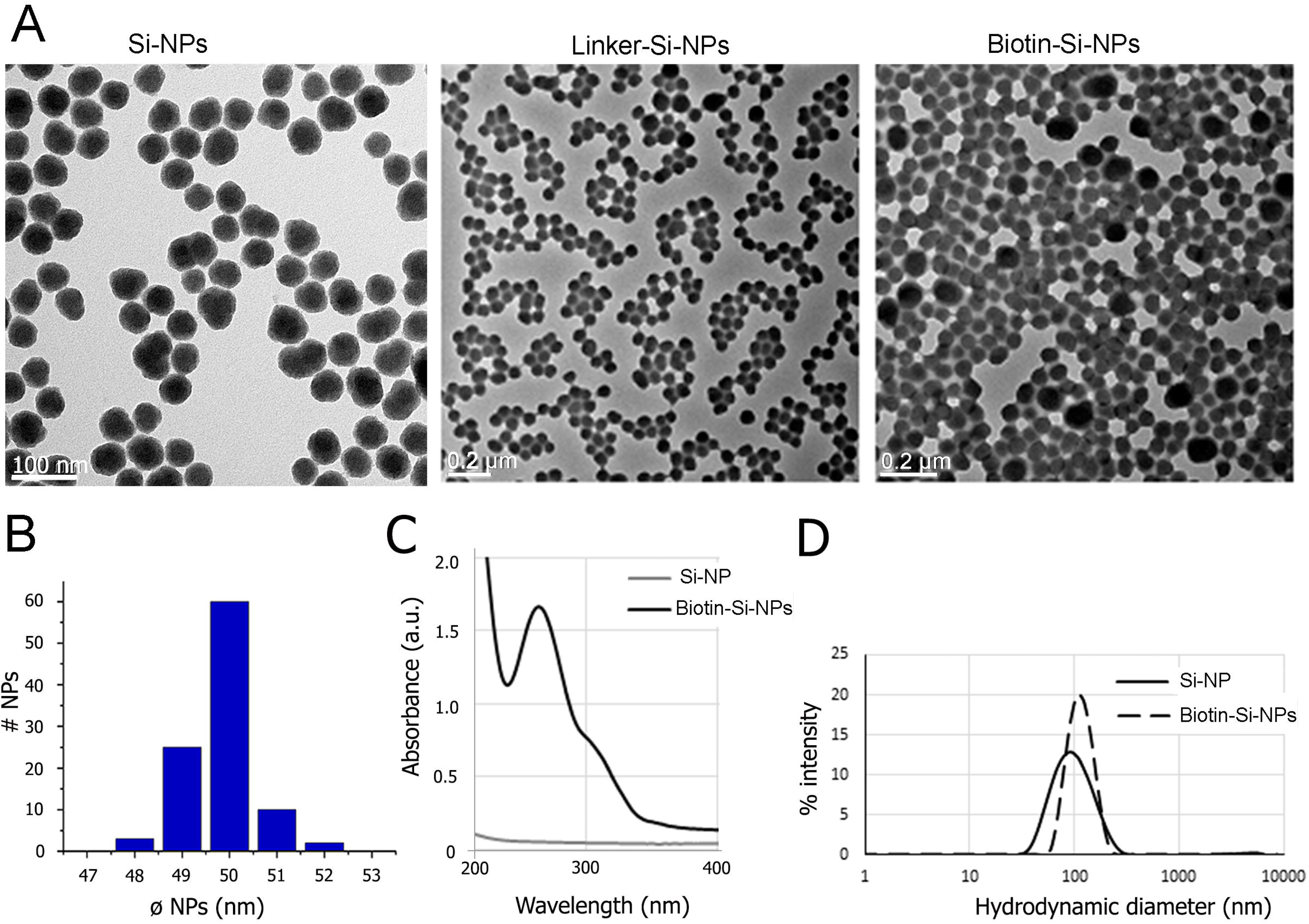
(A) TEM images of Si-NPs before and after functionalization with linker and biotin. (B) Histogram showing the average particle size 50 ± 3 nm before functionalization. (C) Absorbance spectra of Si-NPs (before functionalization) and biotin-Si-NPs (after functionalization). (D) DLS size plots of native and biotin functionalized Si-NPs in water.

A quite narrow size distribution of functionalized NPs was confirmed by DLS measurements. Si-NPs formed monodisperse aqueous solutions of particles with a hydrodynamic diameter (R_H_) of about 90 nm. Conjugation of biotin with the linker moieties shifted R_H_ to about 120 nm (Fig. 1D). This increase is probably related to the hydration layer around biotin units linked to arms bearing negatively charged groups. Biotin-Si-NP solutions were stable at 4°C for at least two months (Supplementary Material, Figs. 1S and 2S).

### 3.2. Concept of biotin-Si-NPs based DNA dot blot

The principle of the DNA dot blot method for *Campylobacter* detection enhanced with biotin-Si-NPs is described in Fig. 2. Initially, the target DNA was immobilized on a porous nylon membrane by UV irradiation to enable crosslinking of DNA to the positively charged surface. Thereafter, the DNA probe CampyP3 labeled with biotin was allowed to hybridize with the target DNA. Hybridization was detected with a streptavidin-HRP conjugate in combination with a chemiluminogenic substrate luminol in the presence of the activator H_2_O_2_. In enhanced detection, the streptavidin-biotin sandwich enabled attachment of biotin-Si-NPs to the DNA probe, and consequently amplification of the detection signal compared to a single biotin.

**Figure 2.**
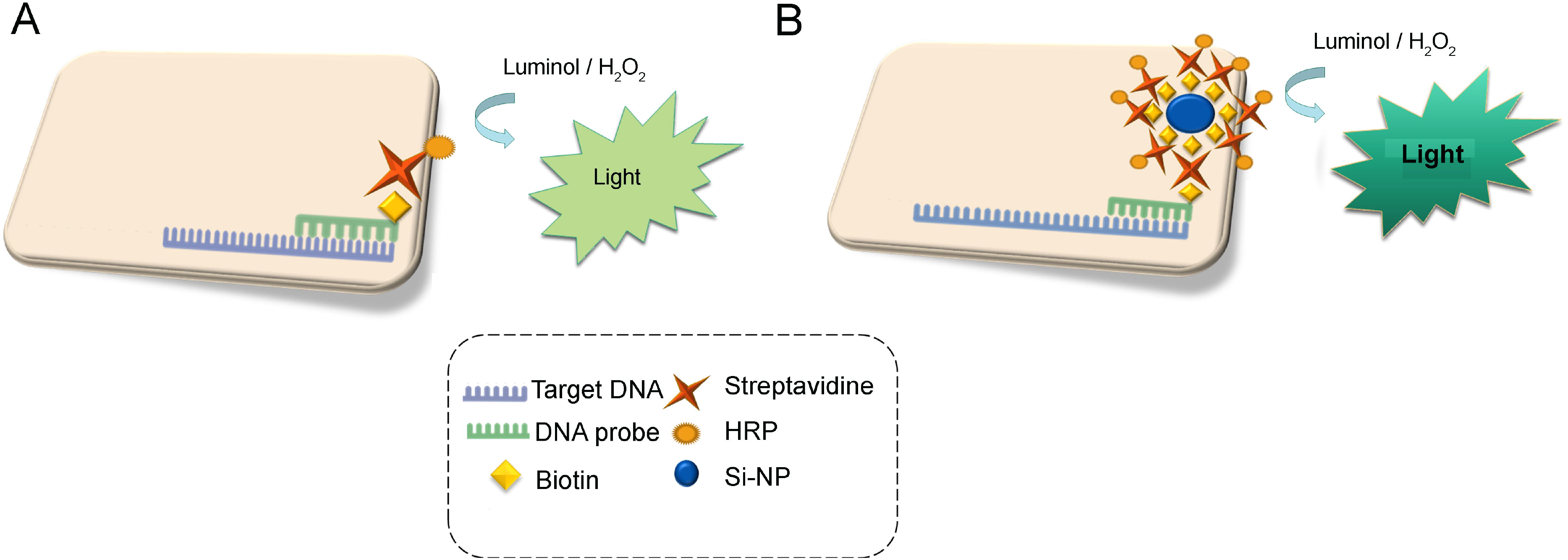
Schematic representation of *Campylobacter* detection based on paper-based DNA hybridization with a complementary biotinylated probe, and a streptavidin-HRP read-out through dot blot (A). The signal was amplified using highly functionalized biotin-Si-NPs instead of a single biotin (B).

### 3.3. Optimization of key parameters and analytical performances of the DNA dot blot sensor

The time and temperature of hybridization of the DNA probe with its target were optimized as they can markedly influence the sensibility and selectivity of a DNA sensor. First, hybridization of DNA was tested at room temperature, 44°C, 55°C and 65°C using complementary and non-complementary short ssDNA targets. A complementary DNA target sequence CP3 (positive control) was detected at all temperatures tested, but only at 65°C no non-specific binding was obtained with negative controls, a truncated target sequence (PR) and *E. coli* sequence (PE) (Fig. 3A). Second, hybridization was studied using 1 ng/μL CP3 at 65°C for different times from 1 h to overnight. The overnight incubation provided the highest level of hybridization and maximal signal intensity.

**Figure 3.**
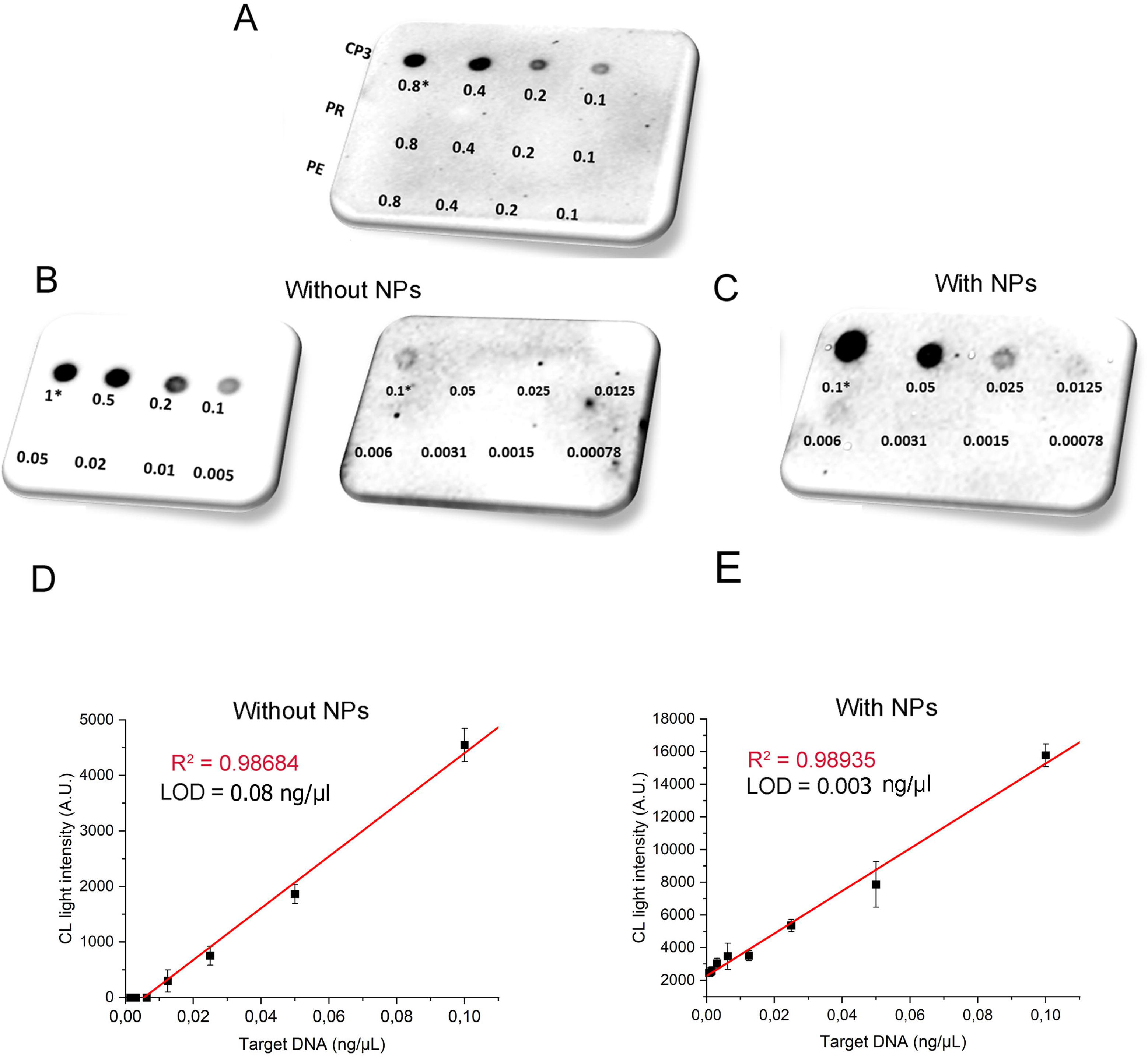
(A) Dot blot detection of *Campylobacter* DNA sequence with biotin-Si-NP enhanced read-out. Note that no signal was obtained with a truncated *Campylobacter* sequence (PR) nor with an *E. coli* control sequence (PE). (B) Conventional dot blot detection of biotin labeled CampyP3 probe using dilution of complementary *Campylobacter* sequence CP3 ranging from 1ng/ μL to 78 fg/μL. (C) Enhanced dot blot detection of CP3 (0.1 ng/μL – 78 fg/μL) using CampyP3 probe. (D, E) Corresponding calibration curves were obtained by plotting the chemiluminescent signal intensity of dots as a function of CP3 template concentrations.

Other parameters that may affect hybridization, such as buffer compositions or the paper support were also tested. The optimization criterion was the best signal-to-background ratio. The nylon membrane allowed for better immobilization of DNA compared to nitrocellulose. The effects of formamide (10, 25, 30, 35 and 50 %), DMSO (0 and 10%) and SDS (0, 1 and 10 %) in the hybridization buffer were tested since these denaturing agents for double strained DNA may prevent nonspecific hybridization. The optimal hybridization buffer contained SDS, and no DMSO or formamide. We determined that addition of 1 % SDS enhanced the intensity of signals obtained and reduced the frequency of unspecific background staining. The effect of SSC concentrations (0.5x, 1x and 2x) in the washing buffer was also investigated. A 3-step washing procedure with two buffers (2xSSC, 0.1 % SDS and 0.5xSSC, 0.1 % SDS) lead to elimination of the background signal while maintaining high light intensity of specific spots.

Detection of 1 ng/μL CP3 under the optimized conditions using various concentrations of streptavidin-HRP and biotin-Si-NPs allowed selection of 25 ng/μL of streptavidin-HRP and 10^6^ nanoparticles/mL for the chemiluminometric reaction. It is worth noting that the chemiluminometric reaction itself does not significantly contribute to background staining as the emission of light arises from the enzymatic reaction without any photonic excitation (Laios et al. 2010).

### 3.4 Calibration curve

Calibration curves were obtained from quantification of chemiluminescent light intensity for different CP3 concentrations, taken from at least three independent dots per concentration. The linear range for the CP3 target was from 1 ng/μL to 0.1 ng/μL (with correlation R^2^=0.98684), and from 6 pg/μL to 0.1 ng/μL (with correlation R^2^=0.98935), for classical and biotin-Si-NP enhanced dot blot techniques, respectively (Fig. 3B-D). The limit of detection (LOD) of 0.08 ng/μL and 0.003 ng/μL for classical and enhanced dot blot, respectively, were calculated using 3 s/m formula, where ‘s’ is the standard deviation of the blank solution and ‘m’ is the slope of the linear calibration graph. Taking into account the molecular weight of the CP3 of 14395.5, the calculated LOD was 5.5 nM and 0.2 nM, for classical and enhanced dot blots, respectively. Consequently, biotin-Si-NPs allowed for almost a 30 fold increase in chemiluminescent signal intensity compared to the classical dot blot. We estimate LOD of about 600 cells considering that one *Campylobacter* cell contains 2 fg of genomic DNA (Pacholewicz et al. 2013). The low LOD obtained suggests that this enhanced dot blot method may be suitable for meat sample testing as the *Campylobacter* infection dose is around 500 cells (2.5 pg DNA).

The proposed biosensor showed comparable or enhanced performance in relation to recently reported DNA sensors for *Campylobacter* detection where the detection limit was 0.5 ng/μL in (Fontanot et al. 2014), 0.37 ng/μL in (Manzano et al. 2015) and 0.09 nM in (Morant-Miñana and Elizalde 2015). The biotin-Si-NP enhanced dot blot reproducibility was estimated to 5 % according to the signal obtained for detection of the same concentration of CP3. Although our time of analysis is not the lowest reported for *Campylobacter* detection (Table 1), our assay is, as far as we know, the first one for the *Campylobacter* detection that has a high sensitivity and does not require the specific bacterial morphology or the background matrix. The colorimetric aptasensor was reported to detect spiral *Campylobacter* cells but not spherical or coccoid ones (Kim et al. 2018); whereas real-time PCR can be inhibited by molecules and ions presented in enrichment broths or meat (Alves et al. 2016).

**Table 1.** Analytical parameters of methods for *Campylobacter spp.* detection.

| Method | LOD | Analysis time | Reference |
| --- | --- | --- | --- |
| Culture-based methods | 1 CFU / 25 g | $\geq 5$ days | (Vizzini et al. 2019) |
| Real-time PCR | $3 \times 10^3$ CFU/mL | 0.5 to 4 h | (Alves et al. 2016) |
| Dot blot | 0.37 ng/μL | 24h | (Manzano et al. 2015) |
| Colorimetric aptasensor | $7.2 \times 10^5$ CFU/mL | 30 min | (Kim et al. 2018) |
| Enhanced dot blot | 3 pg/μL or 600 CFU | 24h | This work |

#### SEM characterization of the biotin-Si-NPs dot blot

SEM was applied to examine different detection steps in order to directly visualize biotin-Si-NPs on the surface of the paper carrying hybridized DNA. The nylon surface before DNA immobilization showed a typical membrane structure, including membrane pores (Figs. 4 and 3S). After *C. jejuni* DNA cross-linking to the nylon, the surface was occupied by a large quantity of ssDNA. Compared to the surface of the bare membrane, the surface of membrane carrying DNA became irregular showing increased morphological heterogeneity. ssDNA molecules seemed to rest horizontally positioned over the surface (Fig. 4, right upper panel). The membrane with hybridized DNA carrying biotin-Si-NPs (final detection step) had thicker fibril structures than the ones with only immobilized ssDNA molecules (Fig. 4, lower panel). Biotin-Si-NPs were easily detected on double strain DNA as small beads of about 50 nm in diameter. It is worth to note, that because the DNA probe matches three points in the *Campylobacter* genome, several biotin-Si-NPs may specifically bind to one DNA. Overall, SEM images confirmed efficient DNA immobilization and hybridization by revealing that the nylon surface became more complex after each stage of the detection process and that hybridized DNA were decorated with nanoparticles of the expected size of 50 nm.

**Figure 4.**
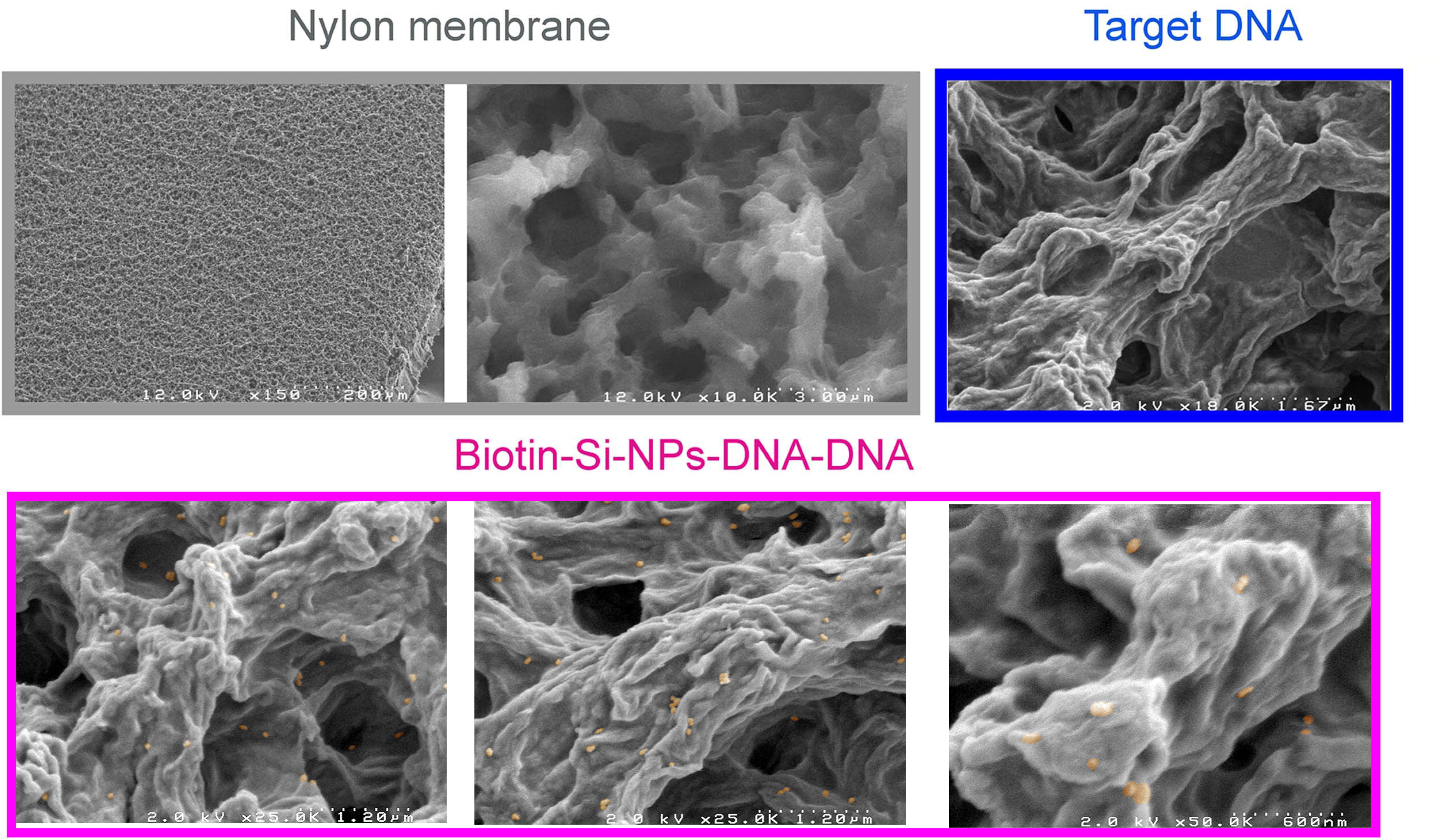
SEM images of the basic nylon membrane, nylon membrane functionalized with a *C. jejuni* DNA, and nylon membrane with CampyP3 hybridized with *C. jejuni* DNA and with biotin-Si-NPs. Note that the size of visualized circular beads of about 50 nm (yellow shading) corresponded to the estimated NP size in Fig 1B.

### 3.4. Specificity and selectivity of detection

To examine the selectivity and sensitivity of the biotin-Si-NPs dot blot, we tested various strains of *Campylobacter* spp. for inclusivity, and 17 other bacterial species and 1 yeast strain for exclusivity (Fig. 5). All strains were cultivated as monocultures and genomic DNA were extracted as explained in the experimental part. Dot blot analyses were performed using 10 ng/μL of non-amplified extracted DNA. Before immobilization, all genomic DNA were denatured for 5 min at 95°C to allow double strain opening. CP3 at 0.1 ng/μL was used as an internal control to normalize signal intensities. The signal ratios for *C. jejuni*, *C. coli*, *C. lari*, and *C. upsaliensis* over CP3 were higher than 1.0, with the membrane images exhibiting obvious dots. Moreover, the intensity obtained for *C. jejuni* and *C. coli* were about 4 times higher than that of the positive control. In contrast, the highest ratio among control DNA and CP3 was only 0.6 for *S. enterica* (background staining). The obtained results highlight the specificity of detection. In addition, the CampyP3 detection probe reacted with DNA of *C. jejuni* and *C. coli* more efficiently than with DNA of other *Campylobacter* spp. tested. Our results demonstrated that the biotin-Si-NPs enhanced dot blot platform is sensitive enough to detect the whole DNA extracted from the most frequent *Campylobacter* spp. and could distinguish between *Campylobacter* and other bacteria. This specificity of the CampyP3 probe suggests that no nonspecific signal will be generated upon testing DNAs of competing total mesophilic flora in chicken meat.

**Figure 5.**
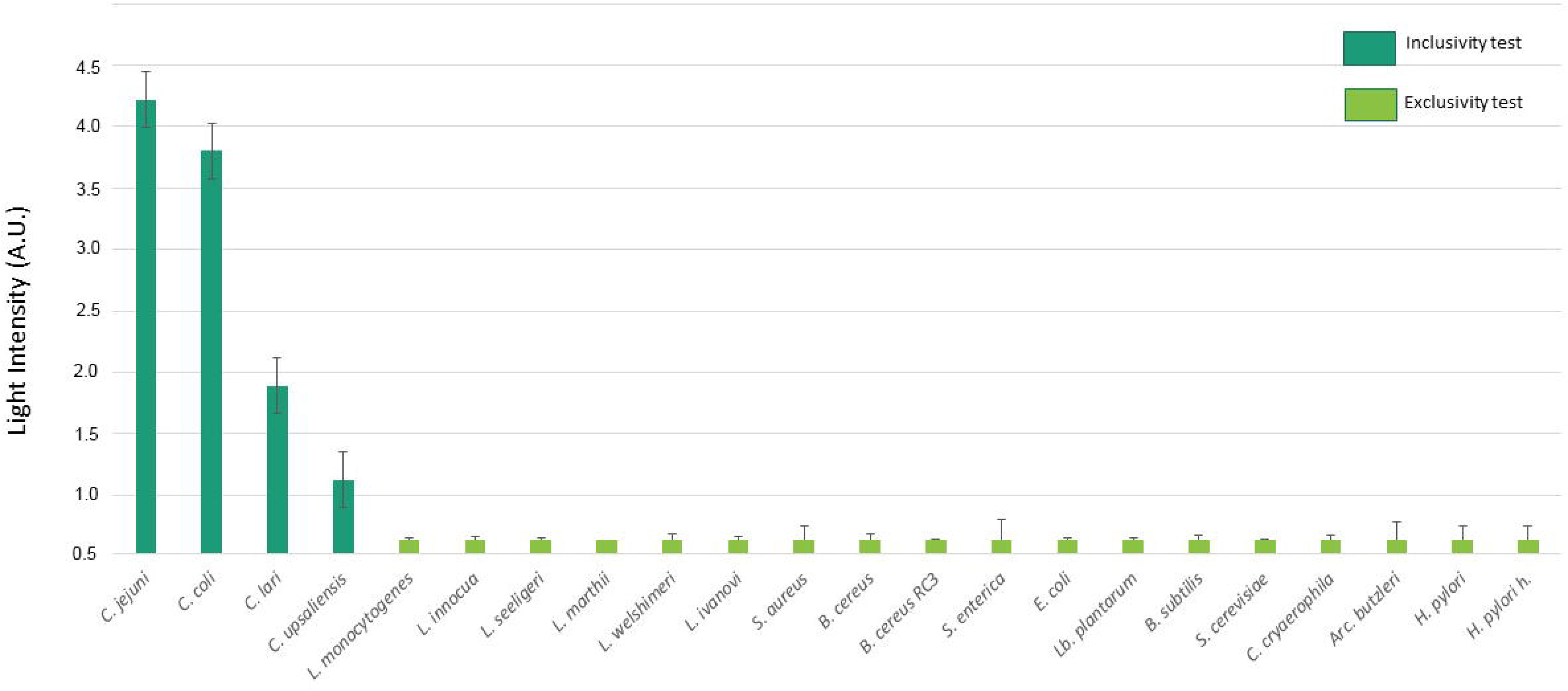
Inclusivity and exclusivity test results of the enhanced dot blot biosensor in pure bacterial cultures. Signal intensity were normalized using an intern control (CP3 template). Error bars represent the standard deviation of the mean from triplicates.

### 3.5. Campylobacter spp. detection in naturally contaminated chicken samples

To demonstrate the capabilities of our developed biosensor for possible real-world applications, we have chosen to target the detection of *Campylobacter* in chicken meat samples. The results of plate count enumeration for background bacteria in five chicken samples are reported in Table 3S. The ISO 10272-1:2006 indicated that two out of five samples were contaminated with *Campylobacter* (Table 4S). Indeed, only isolates from selective media (mCCDA, Skirrow and Columbia agar) of samples C3 and C10 were confirmed for *Campylobacter* by the temperature growth and motility test.

In dot blot analysis, the CP3 template (0.1 ng/μL) was used as a positive control. The dot blot enhanced with biotin-Si-NP successfully detected two contaminated samples, C3 and C10 (Fig. 6). The test achieved a relative specificity of 100 % due to the high specificity of the CampyP3 detection probe. The proposed biotin-Si-NP enhanced dot blot method, thus, specifically detected low levels of naturally present *Campylobacter* after enrichment and could be considered for determination and detection of *Campylobacter* spp. in contaminated chicken meat. Compared to the official ISO 11272: 2006 method, the new DNA dot blot biosensor is less laborious, more cost-effective and time-saving. First, the highly selective DNA detection probe enables selective detection of *Campylobacter* in the presence of meat background bacteria, and thus no culturing on a selective agar is needed. Second, enhancement of the detection signal intensity allows DNA detection without PCR pre-amplification.

**Figure 6.**
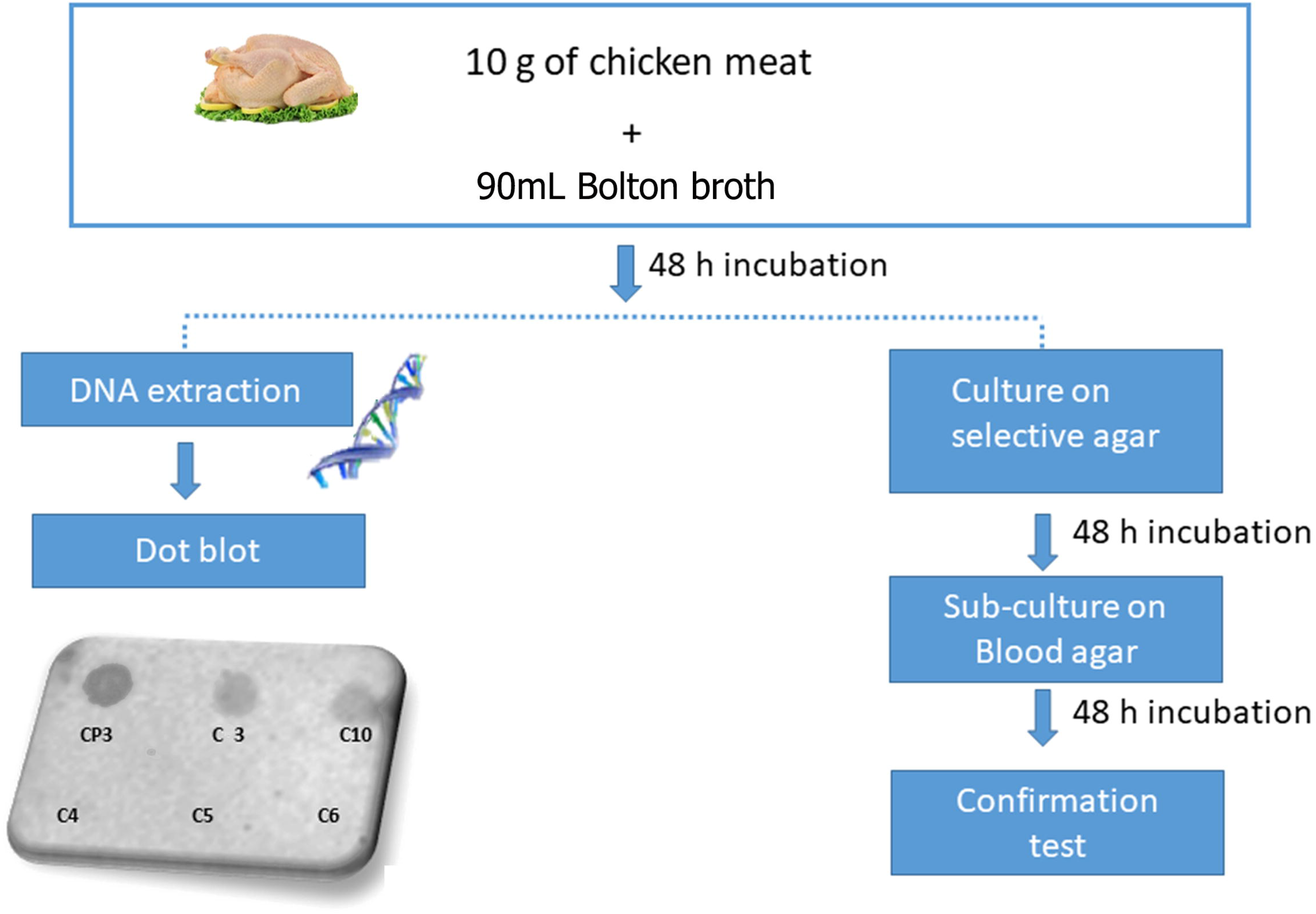
Schematic diagram of *Campylobacter* detection from chicken samples using the enhanced dot blot and the official ISO 10272-1:2006 method. The ISO method involved steps performed to obtain the results given in Table 4. Dot blot membrane shows representative dot blot results obtained with naturally infected chicken samples (C3 and C10) and non-infected chicken samples (C4, C5 and C6). CP3 template sequence (0.1 ng/μL) was used as a positive control.

## Conclusion

This work describes the DNA dot blot method enhanced with biotin-Si-NPs for easy, fast and reliable detection of *Campylobacter* spp. in contaminated chicken meats. The dot blot test is widely used in molecular biology and genetics for detecting target DNA/RNA sequences. This paper-based hybridization can easily analyze multiple samples inexpensively in a high-throughput fashion but has limitation in a low sensitivity. We demonstrated that association of highly functionalized biotin-Si-NPs with the chemiluminescent read-out of the dot blot enhanced the signal about 30 times. Furthermore, biotin-Si-NPs are robust at working temperature and stable in time. The LOD obtained was 3 pg/μL (0.2 nM). Such a low concentration of detected DNA is close to values that can be detected by qPCR (Manzano et al. 2018; Vidic et al. 2019). Hybridization of immobilized genomic DNA with the CampyP3 probe induced positive signals for different *Campylobacter* spp. responsible for human gastroenteritis, while no background staining was observed with control samples. The developed system is a promising tool for fast and cheap screening of poultry samples for the presence of *Campylobacter* since detection is performed on bacterial DNA without a pre-amplification step. Furthermore, our dot blot can be applied on DNA extracted by different extraction methods and from various food matrices, as it is not sensitive to DNA polymerase inhibitors.

The proposed sensitive, miniaturized and multiplex paper-based test is simple to design and could be used by the food industry and regulatory agencies for the detection of other pathogens to monitor food quality. We believe that in the future it could be integrated into a Lab-on-the chip based biosensor that comprises an automatized DNA extraction protocol and a mobile phone camera (Kalligosfyri et al. 2019; Vidic et al. 2019).

## Supporting information

Supplementary information: Highly Sensitive Detection of Campylobacter spp. in Chicken Meat using a Silica Nanoparticle Enhanced Dot Blot DNA Biosenso

## Appendix A. Supplementary material

A list of oligonucleotides and bacterial strains used in this studies, stability of biotin-Si-nanoparticles, as well as the results obtained by the plate count method on chicken samples are available in the supplementary material associated with this article.

## Acknowledgements

The authors thank the Centre Technologique des Microstructures of the Lyon 1 University for providing access to its TEM facilities, and the MIMA2 platform Jouy en Josas for access to electron microscopy equipment (MIMA2, INRAE, 2018. Microscopy and Imaging Facility for Microbes, Animals and Foods, https://doi.org/10.15454/1.5572348210007727E12). JV thanks Maria-Vesna Nikolic (University of Belgrade, Serbia) for English editing. PV acknowledges a doctoral fellowship from the University of Udine, Italy. This research was supported in part by the European Union’s Horizon 2020 research and innovation programme under the Marie Skłodowska–Curie grant agreement No 872662 (IPANEMA), the European Union’s Horizon 2020 research and by the University Paris-Saclay through the Poc in labs 2019 grant agreement No 00003469 (OSCAR).

## Notes

### Competing Interest Statement

P.V, M.M., C.F., C.C, N.R., J.V. declare competing interests. Two application patents have been filed on behalf of INRAE, CNRS and the University of Udine.

### Summary of Updates

This paper is now accepted in Biosensors and Bioelectronics

