## Supplementary information: Highly Sensitive Detection of Campylobacter spp. in Chicken Meat using a Silica Nanoparticle Enhanced Dot Blot DNA Biosenso for "Highly Sensitive Detection of *Campylobacter spp.* in Chicken Meat using a Silica Nanoparticle Enhanced Dot Blot DNA Biosensor"

### 1. List of oligonucleotides used in this study.

**Table 1S.**

| **DNA sequences** | |
|---|---|
| Detection probe (CampyP3 probe) | 5’-[biotin] TAG TGG CGC ACG GGT GAG TAA GGT ATA GTT AAT CTG-3’ |
| Complementary probe (CP3 target) | 5’-CAG ATT AAC TAT ACC TTA CTC ACC CGT GCG CCA CTA-3’ |
| Non-complementary probe PR (rundom) | 5’- CGT GCG CCA CTA CAG ATT ACC TTA AAC TAT CTC ACC-3’ |
| Non-complementary probe PE (*E. coli* target PE) | 5’- GAA AAG CGC GCT GGT GAA AAA AGA GAT GCG CGT CTT-3’ |

**2-*Campylobacter* spp. and control bacterial strains used in this study.**

**Table 2S.**

| **Microorganism** | **Collection code** |
|---|---|
| *Campylobacter jejuni* | DSM[a] 4688 |
| *C. coli* | DSM[a] 24155 |
| *C. lari* | DSM[a] 11375 |
| *C. upsaliensis* | DSM[a] 5365 |
| *C. cryaerophila* | DSM[a] 7289 |
| *Helycobacter pylori* | DSM[a] 7492 |
| *H. pylori* | ICS[b] |
| *Arcobacter butzleri* | DSM[a] 8739 |
| *Listeria monocytogenes* | ATCC[c] 7644 |
| *L. innocua* | DSM[a] 20649 |
| *L. seelligeri* | DSM[a] 20751 |
| *L. marthii* | DSM[a] 23813 |
| *L. welshimeri* | DSM[a] 15452 |
| *L. ivanovii* | DSM[a] 52491 |
| *Staphylococcus aureus* | DIAL[d] |
| *Bacillus cereus* | DSM[a] 4282 |
| *B. cereus* | DIAL[c] RC3 |
| B. *subtilis* | DSM[a] 4181 |
| *Salmonella enterica* | DSM[a] 9378 |
| *Escherichia coli* | DISTAM[e] |
| *Lb. plantarum* | ATCC RAA[b] 793 |
| *Saccharomyces cerevisiae* | ATCC[b] 36024 |

[a]DSM: Deutsche Sammlung von Mikroorganism und Zellkulturen GmbH (Braunschweigh, Germany); [b]ATCC: American Type Culture Collection (Manassas, VA, USA); [b]DIAL : Dipartimento di Scienze e Tecnologie Alimentari (Udine, Italy); [d]DISTAM: Dipartimento di Scienze e Tecnologie Alimentari e Microbiologiche (Milan, Italy); [e]Isolated from Clinical samples (Hospital of Udine, Italy).

### 3- Stability of biotin-Si-nanoparticles

The stability of biotin-Si-nanoparticles over time was tested in samples stocked in water, 50 mM Hepes buffer, pH 7.2, 50 mM Tris buffer containing 5mM $NaN_3$, pH 7.2 and ethanol at 4°C. Nanoparticles functionalized with about 2680 oligonucleotide linkers per NP were at concentration of 5 µM in all samples.

Nanoparticles were stable in both Tris and Hepes buffer for two weeks when stocked at 4°C but started to degrade in pure water. However, in ethanol they were stable for at least two weeks.

It is worth to note, that the nanoparticles were highly stable over 6 months when stored at -20°C.

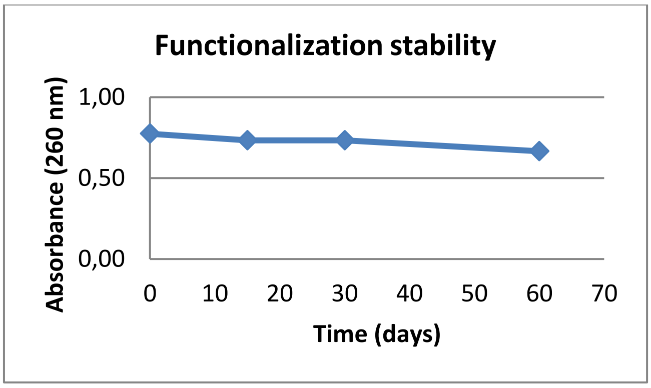


**Figure 1S. Stability of biotin-Si-nanoparticles admixed to pure ethanol and stored at 4 °C was followed over time by measuring the linker absorbance at 260 nm.**

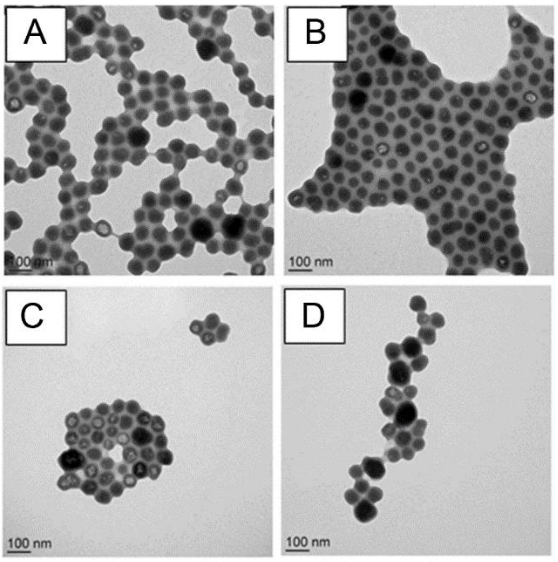


**Figure 2S. Stability of biotin-Si-nanoparticles admixed to pure ethanol and stored at 4 °C was followed over time by ultrastructural visualization using scanning electronic microscope at (A) day 1 (B) day 15, (C) day 30 and (D) day 60.**

**3- SEM visualization of different steps involved in detection**

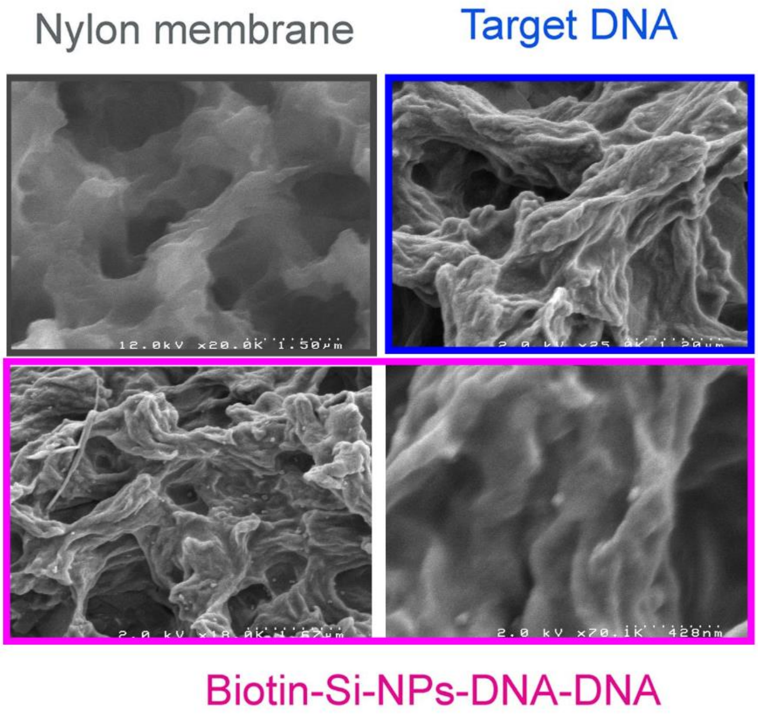


**Figure 3S. SEM images of basic nylon membrane, nylon membrane functionalized with a *C. jejuni* DNA, and nylon membrane with CampyP3 hybridized with *C. jejuni* DNA and revealed with biotin-Si-NPs.**

**4-Plate count data of chicken (C) samples expressed as log of Colony Formin Unit (CFU/g).**

**Table 3S.**

| Sample | Total viable count | Enterobacteriaceae | Coliforms | *E. coli* | Yeasts | Molds |
|---|---|---|---|---|---|---|
| C3 | 5.3 | 3 | 1.1 | 2.9 | 2.9 | 0.7[1] |
| C4 | 6.7 | 4.1 | 3.5 | 3.7 | 5 | 0.7[1] |
| C5 | 7.2 | 4.3 | 3.8 | 3.1 | 5.8 | 0.7[1] |
| C6 | 5.8 | 3.2 | 3.4 | 2.9 | 4.8 | 0.7[1] |
| C10 | 4.1 | 3.1 | 2.9 | 1.9 | 2.2 | 0.7[1] |

[1], under the limit of detection

**Table 5S. The presence /absence of *Campylobacter* spp. in chicken (C) samples assessed with the ISO 10272-1:2006 protocol.**

| Sample | mCCDA* | SKR§ | CBA° | Confirmation Absence/presence 41.5°C | 25°C | Oxidase | Motility | Results |
|---|---|---|---|---|---|---|---|---|
| C3 | + | - | + | - | - | + | + | + |
| C4 | + | - | + | + | + | + | - | - |
| C5 | + | - | + | + | + | + | - | - |
| C6 | + | - | + | + | + | + | - | - |
| C10 | + | - | + | - | - | + | + | + |

*mCCDA, modified charcoal and deoxycholate agar selective medium; §SKR, Skirrow selective medium; §CBA, Columbia blood agar.
